# Detecting and Quantifying Networks of Biological Kinship via Exponential Family Random Graph Models

**DOI:** 10.1101/2025.06.25.661441

**Authors:** Adam B. Rohrlach, Guido Alberto Gnecchi-Ruscone, Zuzana Hofmanová, Matthew Roughan, Wolfgang Haak, Jonathan Tuke

## Abstract

Genetic relatedness between ancient humans can help to identify close and distant connections between groups and populations, uncovering signatures of demographic histories such as identifying mating networks or long-range migration. Critical to researchers are the characteristics that connected individuals, or groups of individuals, share, and how these characteristics interact and are correlated. Here we use Exponential Random Graph models as a method to explore demographic and contextual parameters that may help to explain the significant drivers of the topology of mating networks, as well as to quantify their effects. We show through simulations that model selection and coefficient estimators facilitate the exploration of such networks, and apply the method to individuals from a collection of Avar-associated cemeteries from the Carpathian Basin dating to the 6^th^ to the 9^th^ centuries CE.

## Introduction

Past and present human populations actively and passively engaged in social, commercial and cultural practices that led to the formation of mating networks(1–6). These networks, which can be thought of as massive family pedigrees, were distributed across time and space. The processes that led to the formation of these networks, and which societal norms mediated their shape and topology, are largely unknown, and it is unclear how to leverage the evidence in the observable archaeological record to fill this gap.

Recent developments in ancient DNA (aDNA) analyses and ever-growing sample sizes (both per site and in the field of archaeogenetics overall) have allowed researchers to estimate snapshots of these networks of close and distant relatedness from both low- and high-coverage sequence data(7–10). These methods have been applied to whole-cemetery analyses of single sites, as well as intra- and cross-regional studies, to reconstruct deep pedigrees(11–16). However, these methods can also uncover inter-regional connectedness due to contact, trade and exchange, individual mobility, and migration.

Studies have so far focused on single statistics calculated on networks without taking into account potential network-wide dependency structures, and answer only limited hypotheses about single parameters, without quantifying these effects or considering potentially interacting factors(12, 15).

Exponential Random Graph models (ERGMs) are a powerful tool that have been used to analyse both sparse and dense networks with interacting covariate effects and a wide range of complex dependencies(17). ERGMs have been used to explain the structure of social networks, owing to their ability to detect significant predictors even when the overall model explains only a relatively low percentage of the variability in the network, such as might be expected in mating networks(18). Hence, ERGMs are a sensible tool to apply to networks constructed from genetic relatedness, but that also contain genetic, archaeological and anthropological variables measured on the individuals. Importantly, using ERGMs, we can quantify the impact of variables on the connectedness of individuals, and yield measures of fold-increase in the probability of “types” of individuals sharing a genetic relationship.

To highlight the usefulness of ERGMs to efficiently detect variables of importance and to quantify their impact, we use simulated networks and Bayesian Information Criterion based model selection to show that we can reliably recover the correct, and hence best-fitting, models. We then explore how sample size and the relative effect size of variables impact the performance of ERGMs to give researchers a baseline for when ERGMs can be reliably applied. Finally, we use ERGMs to analyse an empirical data set from four Avar-associated cemeteries from the Carpathian Basin to show how ERGMs can be used to combine archaeogenetic and archaeological data into one cohesive analysis.

## Methods

### A. The statistical model

Consider an undirected network *G* with a set of nodes *V* (*G*) = {1, 2, …, *n*} where *i* represents the *i*^th^ individual (with |*V* | = *n* total individuals in the study) and a set of edges *E*(*G*) which represent pairs of individuals with a given type of relationship, such as sufficient shared IBD (identical by descent). We can expand this approach to use a directed network if the relationships are not symmetric (*e*.*g*.,pedigree relationships) but we do not consider directed graphs here.

It is common to form such a network starting with a measure *e*_*ij*_ of the connectedness of individuals *i* and *j* formed from genetic or other data, where 0 indicates no measured connection, and a high value indicates a strong relationship. We then construct the network through thresholding, *i*.*e*.,we define the network adjacency matrix **Y** = [*Y*_*ij*_], where

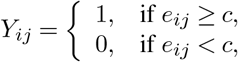

where *c* is a cutoff that determines a significant connection. An edge (*i, j*) exists in *E*(*G*) if and only if *Y*_*ij*_ = 1.

The choice of cutoff might be determined through a statistical model of the data-measurement process, or by calibration to some ‘ground-truth’ data, for instance, obtained through *in silico* simulations. We then treat our network as a random sub-sample of the complete network, the sub-sample being determined via measurement noise, and the presence or absence of measured individuals.

Additionally, covariates are recorded for individuals and edges, denoted **V** and **W**, respectively. For example, node covariates **V** record demographic parameters about each individual such as

- the genetic sex,
- age at death or,
- burial context.

The edge covariates **W** record pairwise shared measures, such as

- the distance between burials or
- the difference in mean ^14^C date.

Of interest is the probability of an edge existing in the network, denoted

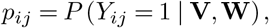

conditional on these covariates. Moreover, we assume locality in the sense that the probability of connection between individuals *i* and *j* depends only on the covariates of these individuals, *i*.*e*.,**v**_*i*_, **v**_*i*_ and **w**_*ij*_. We can further construct synthetic edge covariates **z**_*ij*_ from pairs of node attributes, *e*.*g*.,we might consider *homophily* measures such as *z*_*ij*_ = *I*(*v*_*i*_ = *v*_*j*_), where *I*(·) is the indicator function (which takes the value 1, when its argument is true, and 0 when false).

As in logistic regression, we work in the log-odds space and instead look to make inferences about log(*p*_*ij*_*/*(1− *p*_*ij*_)). We perform a network regression to model the log-odds of any two individuals being connected given the measured covariates, *i*.*e*.,

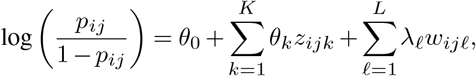

where *θ*_*k*_ and *λ*_*/l*_ are the coefficients for the *K* node and *L* edge variables *z*_*ijk*_ and *w*_*ij/l*_, respectively. For simplicity we restrict the following examples to node attributes, and hence assume *L* = 0.

While this approach has many similarities to logistic regression, there are some critical differences. First, the assumption of independence for observations: in the network case, connectedness of nodes can not be assumed to be independent. To see this, consider that if individuals *i* and *j*, and individuals *i* and *k* are closely genetically related, then it is likely that individuals *j* and *k* are also closely related.

Second, the nature by which nodes manifest relationships can be varied—consider the example of individuals buried at three sites: A, B and C, with genetic sexes “Sex 1” and “Sex 2” (for a full discussion see Section A).

It might be that simple homophily (nodematch) is the primary factor determining connectedness, that is, a pair of nodes that *share* an attribute are more likely to be connected. An example of this may be that individuals that share the same burial site are more likely to share a genetic connection.

Simple homophily requires that the increase in connectedness probability is the same for each site. An alternative is differential homophily (differential sitematch), *i*.*e*.,connectedness is still more likely for individuals that share the same attribute, but the value of the attribute (that they share) is also important. For example, it may be the case that individuals at different sites are more likely to share a genetic connection, but that site A was *more* interconnected than sites B or C, and hence two individuals buried at site A are more likely to be connected than two individuals buried at site B (or C).

Finally, we consider the idea of attribute mixing (nodemix). In this case, it no longer matters whether the attributes match, but the combination of the two attributes is of importance. An example of this could be that (as in the previous example) site A is more interconnected, but sites B and C are close to each other. Hence, while individuals within sites are quite likely to be related, and more so for site A, individuals between sites B and C are also more likely to share a genetic connection than between A and B (or A and C).

From a statistical modelling perspective, these are increasingly complex models, *i*.*e*.,they require additional parameters to be estimated. See Section **??** for a list of the models, but for the moment note that in our examples the first requires one parameter (the degree of homophily), the second requires three (one homophily parameter for each region) and the third requires up to six parameters to consider all pairs of regions.

### B. Model selection

Model selection is the process of choosing which factors (covariates) are important in determining the model for connection probabilities, *i*.*e*.,which *z*_*ijk*_ terms are best included in the model. A typical problem is that more complicated (more highly parameterised models) will often fit the data more exactly, but without generalizability, *i*.*e*.,without the ability to extrapolate to datasets other than the one measured. Such over-fitting is highly undesirable and usually avoided using one of several *Information Criteria*. Here we use the Bayesian Information Criterion (BIC)(19). BIC prevents over-fitting by introducing a penalty term for additional model parameters. We use BIC, as opposed to alternatives such as Akaike’s Information Criteria, because (i) it is more conservative, and (ii) it is more naturally matched to the type of models being used here(20). For automated model selection we take the model with the smallest BIC, however when interpreting the BIC, it is important to consider the scale of the improvement in BIC for more complex models: a change in BIC of zero to two units is considered “not worth mentioning,” and hence models within 2.0 of each other are often considered equally valid(20). We also compare the performance of BIC to AIC and a p-value-based approach to show that BIC outperforms all other model selections methods.

### C. Simulation performance and power analyses

To test the performance of ERGMs to be able to identify the correct model we simulated 100 realisations of each model (see Section B), yielding a total of 700 simulations. We then applied the BIC model selection criteria, and recorded the best fitting model for each simulation. From this, we are able to calculate a range of performance statistics, including accuracy and specificity.

We were also interested in calculating a lower cut-off for how impactful variables must be on the network to be detected, dependent on the number of vertices in the network. It should be noted that the number of observed edges is of most importance, but as we kept the expected number of edges roughly constant, variable effect and the number of edges are highly correlated. We simulated networks from the site match model, with the number of nodes |**V**|∈ **{**50, 100, …, 1000}, and with values of *θ*_0_ = −4 and *θ*_1_ ∈{0, 0.05, …, 1}. We use a simple p-value cut-off of *α* = 0.05 for coefficient significance. For each value of |**V**|, we are then able to find the minimum value of *θ*_1_ such that the standard 80% power is achieved(21).

### D. Network clustering and centrality analyses

To investigate structure within the unweighted Avar network we performed Louvain clustering on a network with edges weighted by the total amount of shared IBD, and calculated degree and betweenness centrality using the *igraph* package.

## Results

### E. Performance of ERGMs on simulated data

To evaluate the accuracy of model selection, and hence variable selection, we simulated 100 realisations for each of the seven possible models on networks with *n* = 300 nodes, and parameters values as described in Section **??**. For each realisation we used the minimum BIC value to assign the best possible model. Interestingly, no single realisation was incorrectly assigned to the true model. We also found that BIC outperformed using AIC or the p-values associated with the coefficients of the model for model selection (see Section D). Following this, we were interested in assessing the statistical power of ERGMs when performing model selection using BIC. To start with, we simulated realisations of the site match model (see Section B.2) with varying values of *θ*_1_. We let *θ*_1_ take values from zero to one, in steps of 0.05, which equates to a fold-increase in probability of between one and 2.5 (see Figure 2. We find that when *θ*_1_ = 0, power is the expected 0.05. We also find that for *θ*_1_ ≥ 0.35 (approximately 1.41-fold increase), power is one. Finally, we find that we achieve the standard 80% for *θ*_1_ ≥ 0.25, which is equivalent to a fold-increase in probability of only 1.27, meaning that even subtle variables can be detected using BIC.

**Fig. 1.**
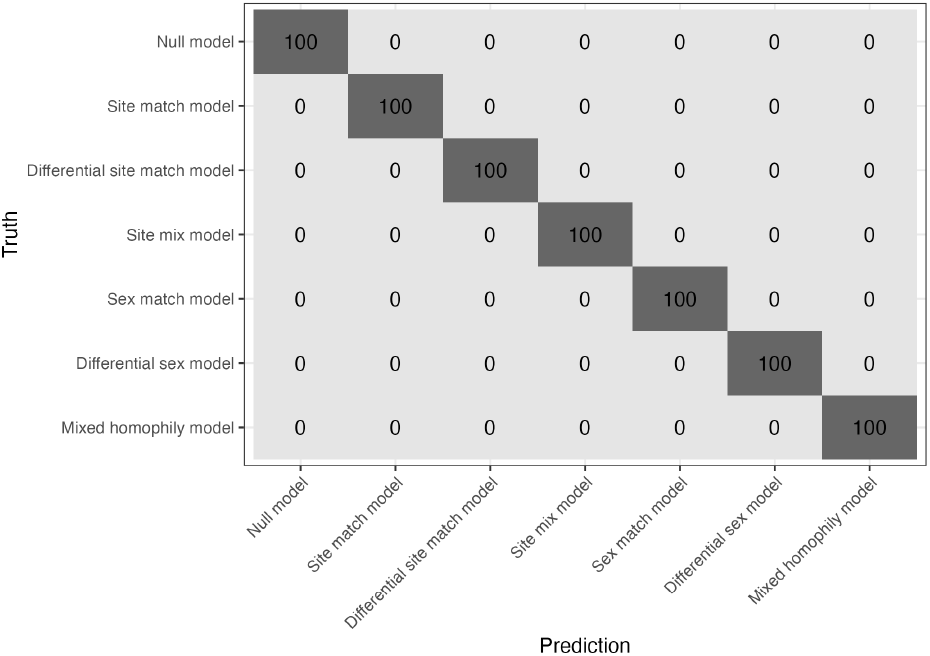
A confusion matrix for the accuracy of model selection via BIC.

**Fig. 2.**
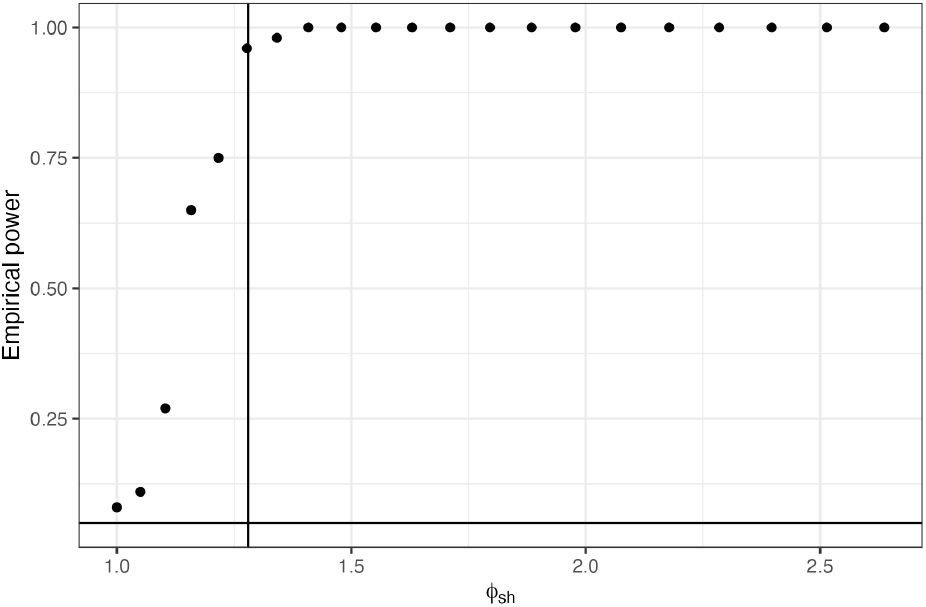
Empirical power calculations for the sex match model. The x-axis indicates the fold-increase in probability if the sex matches for two vertices, and y-axis gives the value of the empirical power.

We tested whether the power to detect these variables may be affected by uneven sampling. To assess this, we repeated the above experiment, but this time simulated that 25% of nodes are “Sex 1”, and that the remaining 75% are “Sex 2”. We find that power is slightly decreased in the unequal case for smaller values of *θ*_1_ (see Figure 3). However, 80% is still achieved relatively quickly at *θ*_1_ = 0.3 (approximately 1.34-fold increase), and that 100% power is achieved quickly after this.

**Fig. 3.**
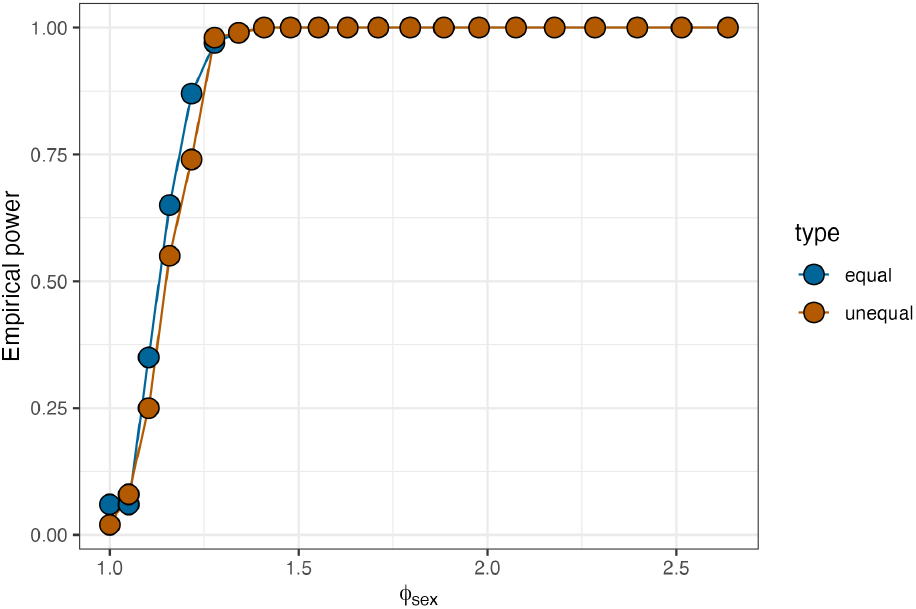
Empirical power calculations for the sex match model. The x-axis indicates the fold-increase in probability if the sex matches for two vertices, and y-axis gives the value of the empirical power. Blue points indicate equal numbers of each sex (50% each), and orange indicates unequal numbers of each sex (25% and 75%).

Finally, we were interested in how the number of vertices in the network affected the power of BIC to correctly perform model selection. We note that it is not technically the number of vertices in the network, but rather the number of edges that will change predictive power. However, we keep this proportion relatively constant and instead vary the number of vertices. We simulated the site match model with the same values of *θ*_1_ again, but this time for networks with between 50 and 1,000 vertices, in steps of 50. For each number of vertices, we returned the minimum value of *θ*_1_ that achieved 80% empirical power (see Figure 4).

**Fig. 4.**
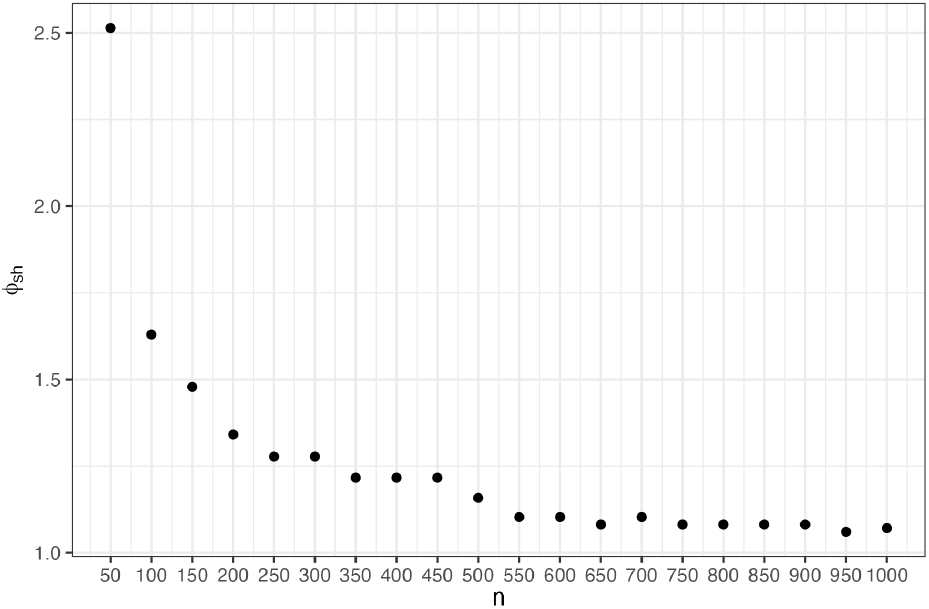
Empirical power calculations for the sex match model. The x-axis indicates number of vertices in the network. The y-axis gives the minimum value of the fold-increase in probability if the sex matches for two vertices, that achieves an empirical power of 80%.

**Fig. 5.**
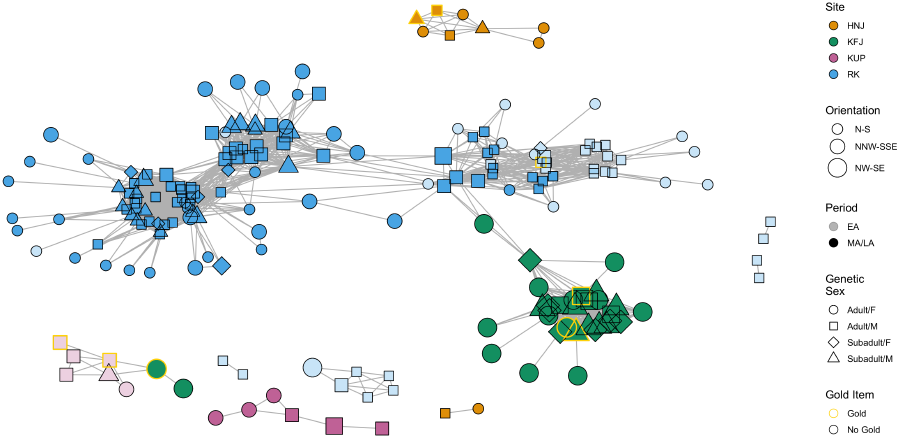
The Avar network connecting individuals from Hajdúnánás-Fürj-halom-járás (HNJ, orange), Kunszállás-Fülöpjakab (KFJ, green), Kunpeszér-Felső peszéri út (KUP, purple) and Rákóczifalva Bagi-földek 8 (RK, blue). Shapes indicates the genetic sex and age of the individuals. Edges exist if two individuals share at least two blocks of IBD of length of at least 12cM, and one block of at least 16cM.

Unsurprisingly, we find that as the number of vertices, and hence edges, increases, that the minimum fold-increase in probability decreases. This is expected, as more edges result in more information, and hence more statistical power. We also see that ERGMs are capable of reliably detecting even very subtle variables, with fold-increases of less than 1.5 for networks with just 150 vertices, tending to a value of approximately 1.07-fold as the number of vertices becomes large.

### F. Analysis of empirical data

Finally, we reanalysed the networks presented in Gnecchi-Ruscone *et al*. 2024(15). The networks represent individuals sampled from four nearby sites, associated with the early to late Avar period in Hungary (567-822 AD). Using measures of centrality, the authors found that the sub-networks connecting genetically male individuals were significantly more dense than either the subnetwork connecting genetically female individuals, or the full network(15). The network consisted of 237 individuals, connected across sites, for which several variables were recorded including genetic sex, age (adult/subadult), body orientation, and whether they were buried with (a) a common item: an iron buckle, or (b) something rare and valuable: a golden item (or both).

Variables that we might expect would lead to higher probabilities of genetic relatedness (due to geographical and temporal closeness) are retained as significant: site and period (see Table 1). Since differential node match worked best for period, it is then the fact that individuals born during the Early Avar period and Middle/Late Avar period are more likely to be related to individuals from the same time period (7.24- and 9.39-fold, respectively). Individuals buried at the same sites are also more likely to be related, and this differs from site to site as some sites are “more inter-related”. Specifically, HNJ (656-fold), KFJ (812-fold) and KUP (580-fold) are more inter-related than RK (166-fold). This discrepancy can also be seen when performing network clustering (so-called community detection) and inspecting the connectedness of the network using centrality measures (see Section E.2). The three visible clusters in RK can be separated into a total five communities, indicating relatively low within-site connectivity. Similarly, KUP is split into two clusters. We observed that these two KUP clusters can be separated by period.

**Table 1.** Summary of best-fitting ERGM for the empirical Avar data. The base case for comparison for the Age/Sex variable was Adult F/Adult F. Coefficients that are not significant are in italics. match = Node Match, Diff = Differential Node Match and Mix= Node Mix.

| Variable | Level | Fold | P-value | Type |
| --- | --- | --- | --- | --- |
| Site | HNJ | 656.00 | $< 10^{-6}$ | Diff |
| Site | KFJ | 811.58 | $< 10^{-6}$ | Diff |
| Site | KUP | 580.44 | $< 10^{-6}$ | Diff |
| Site | RK | 165.60 | $< 10^{-6}$ | Diff |
| Period | Early | 7.24 | $< 10^{-6}$ | Diff |
| Period | Middle/Late | 9.39 | $< 10^{-6}$ | Match |
| Sex/Age | Adult F/Adult F | 0.324 | $< 10^{-6}$ | Mix |
| Sex/Age | Adult M/Adult M | 4.19 | $< 10^{-6}$ | Mix |
| Sex/Age | Adult F/Subadult F | 0.94 | 0.753 | Mix |
| Sex/Age | Adult M/Subadult F | 4.38 | $< 10^{-6}$ | Mix |
| Sex/Age | Subadult F/Subadult F | 2.85 | 0.009 | Mix |
| Sex/Age | Adult F/Subadult M | 1.16 | 0.274 | Mix |
| Sex/Age | Adult M/Subadult M | 4.56 | $< 10^{-6}$ | Mix |
| Sex/Age | Subadult F/Subadult M | 5.59 | $< 10^{-6}$ | Mix |
| Sex/Age | Subadult M/Subadult M | 6.93 | $< 10^{-6}$ | Mix |
| Orientation | Orientation Match | 2.95 | $< 10^{-6}$ | Match |

Interestingly, the age and sex of an individual was the only variable for which the more complex node-mix was selected (see Table 1). We observe that, compared to an adult male and an adult female, two adult males are far more likely to be related (4.19-fold) and that two adult females are far less likely to be related (0.32-fold). This might indicate a social practice of patrilocality and/or female exogamy: where female individuals leave the place of their birth to find a mate and live near the family of their male partner. We also observe that the only other factors that lead to no change in probability compared to adult male/adult female pairs also involves adult females: adult female/ subadult female (*Z* = − 0.315) and adult female/subadult male (*Z* = 1.09). This suggests that when female individuals left their place of birth to join a mate after they reached some age of maturity, that they left behind both living brothers, and siblings who had died before this age was reached.

Variables from archaeological contexts were also included. Both burial item measures (Gold items or Iron Buckles) were not significant. This indicates that these items were either “randomly” given to individuals (with respect to genetic relatedness), or were correlated with other variables (such as period or site), and hence not statistically meaningful. However, orientation was significant, and individuals with matching burial orientation had an approximately 2.95-fold increase in relatedness. This finding supports the observation of Gnecchi-Ruscone *et al*. that different funerary customs are associated with different sub-groups within the overall pedigree, an association between archaeological context and fine-scale genetic affinity that our method identified.

## Conclusions

ERGMs are a powerful method for performing regression on network connectedness, allowing researchers to combine multifaceted variables, from genetics, archaeology, anthropology and other sources, into one analysis. The simple output from these regression models allows researchers to identify variables of importance and to quantify their impact on the network, allowing for intuitive, real-world interpretations of the final model.

We have shown that using the Bayesian Information criterion is the best method for model selection allowing for the comparison of competing models of varying complexity. By investigating the limits of BIC to detect variables of interest with varying effects, it can be seen that ERGMs can detect even very subtle variables, and report the resolution of the statistical power for varying network sizes.

The utility of ERGMs in this setting was demonstrated by reanalysing the Avar network reported by Gnecchi-Ruscone *et al*.. We were able to recreate the key result by showing that genetic sex was a significant predictor for network connectedness. We were also able to show this while accounting for site of origin and time period, both which should affect connectedness, but which could have been correlated with sex due to sampling bias. Further, we were able to show that the age at death was also a significant predictor, further reinforcing the finding of a patrilineal kinship system, in which patrilocality and female exogamy were the norm. We also report a new finding where body orientation was a significant predictor of relatedness.

Overall, as relatedness networks become more common in archaeogenetic studies, we present ERGMs as a proven method for understanding their structure. As in all studies of the human past, it is critical that multiple sources of information are co-analysed, so that the most robust interpretations of the data can be found. ERGMs inherently encourage this approach to interdisciplinary analysis in a statistically rigorous and familiar framework.

## Funding

ABR was supported by the Max Planck Society. This research was funded by the European Research Council under the European Union’s Horizon 2020 research and innovation programme under grant agreement numbers 101141408-ROAMANCE and 856453-HistoGenes.

## Acknowledgements

The authors wish to thank Prof. Johannes Krause for his support in this research, and Dr. Harald Ringbauer, Professor Nigel Bean and Dr. Vincent Braunack-Mayer for enlightening and instructive conversations.

## Supplementary Note 1: Supplementary Notes

### A. Interpretation of coefficients

We give these examples below in model which *only* use site as a predictor for genetic relatedness, where *x*_*i*_ is the variable which indicates the site at which individual *i* was buried.

1. **Null Model**: site location does not affect the probabilities of two individuals being connected, and hence connectedness is simply random.

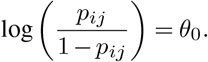 Note then that *θ*_0_ is the log-odds of two random individuals sharing a genetic connection.
2. **Homophily**: All that matters is whether the individuals are from the same site, i.e

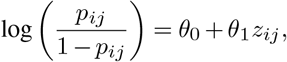

where

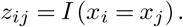 Here, *θ*_0_ is the log-odds that two individuals from *different* sites share a genetic connection, and *θ*_1_ is the amount that the log-odds increase (or decrease) when individuals are buried at the same site.
3. **Differential Homophily**: Whether two individuals are buried at the same site, and which site this is, is of importance, i.e.

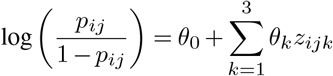

where

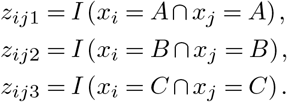

Hence, again *θ*_0_ is the log-odds that two individuals from *different* sites share a genetic connection, and *θ*_1_, *θ*_2_ and *θ*_3_ represent the amount the log-odds increase (or decrease) when individuals are both buried at site A, B or C, respectively.
4. **Attribute Mixing**: the combination of the sites that the two individuals are buried at is of importance.

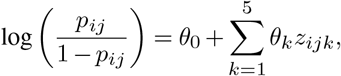

where

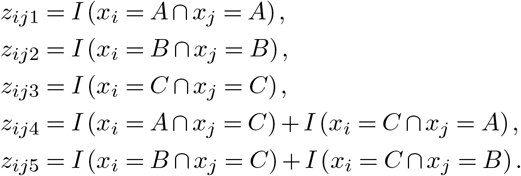

Hence, *θ*_0_ is the log-odds of two random individuals buried at sites A and B, and the *θ*_*k*_ give the increase (or decrease) in the log-odds if the individuals are both buried at site A, B or C, or at sites A and C, or sites B and C, respectively.

### B. Simulated model descriptions

To test the ability of ERGMs to differentiate between different and known model types, we simulated 100 independent realisations of each of the following models. Overall, we allowed nodes to have one of two attributes. First, genetic sex can take one of two values: female (F) or male (M). Second, archaeological site (referred to as “site”) can take one of three values: A, B or C.

#### B.1 The null model

ℳ_0_. The null model is a model in which no variable explains the connections between nodes. The genetic sex and the site are not predictors of when two individuals are connected.

Here we set *θ*_0_ = − 4, and all other values of *θ*_*i*_ = 0. This corresponds to a probability of approximately 0.0183 of any two random individuals being related. This is the base probability used in all models.

#### B.2 The site match model

ℳ_1_. The site match model makes any two individuals equally more likely to share a connection, no matter which site they are attributed to. This causes the individuals to cluster by site, and those clusters look equally densely connected.

Here, *θ*_1_ = *θ*_2_ = *θ*_3_ = 3, corresponding to 14.95-fold increase in the probability of individuals being connected if they share the same burial site.

#### B.3 The differential site match model

ℳ_2_. The differential site match model makes two individuals more likely to share a connection if they are buried at the same site, but not equally, and the increase depends on *which* site they are both buried at. This causes the individuals to cluster by site, and for those clusters to look more densely connected depending on the scale of the increase.

**Fig. 6.**
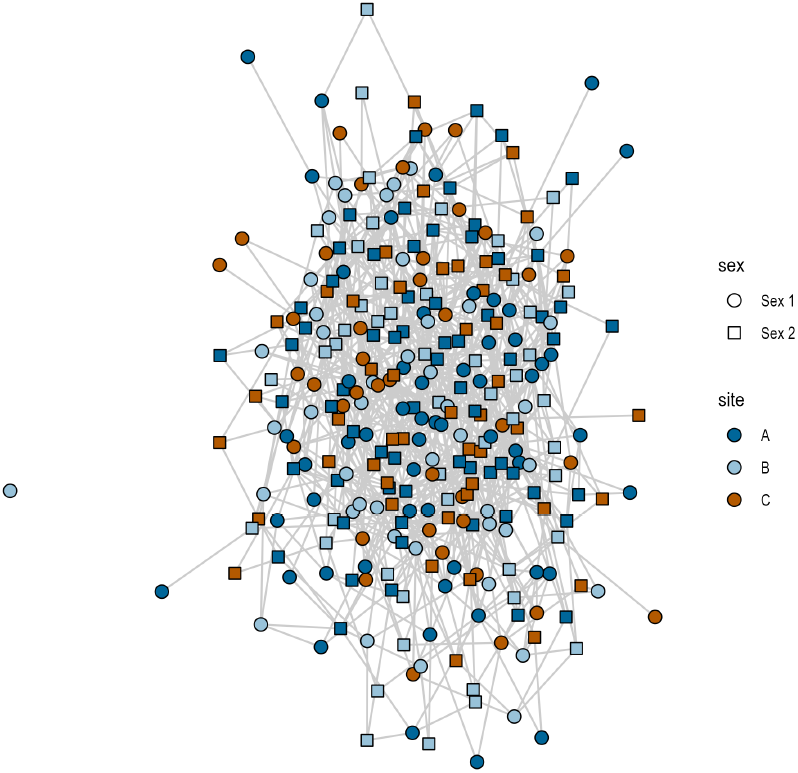
A single realisation of the null model, ℳ_0_.

**Fig. 7.**
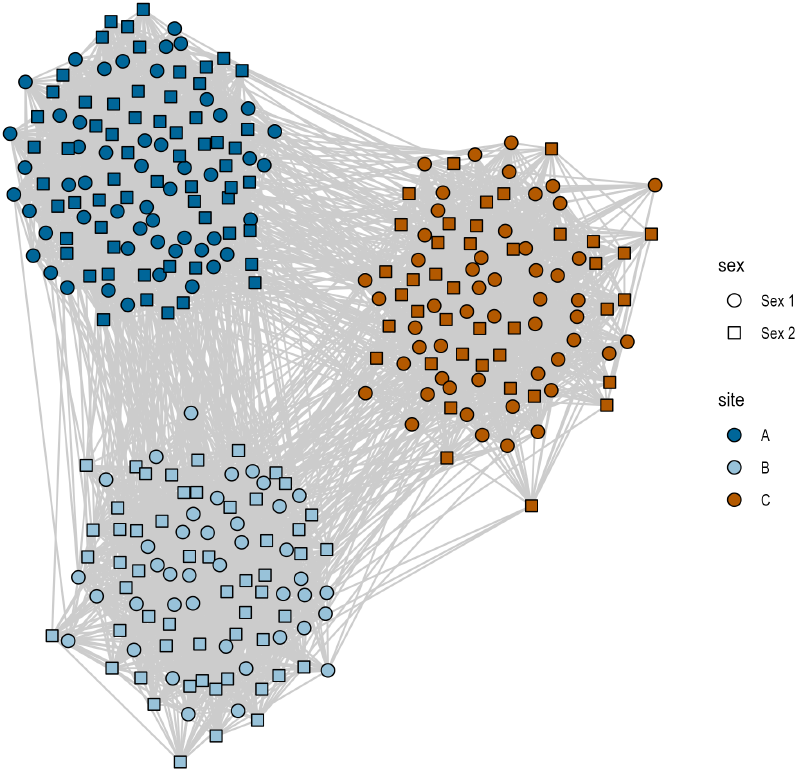
A single realisation of the site match model, ℳ_1_.

**Fig. 8.**
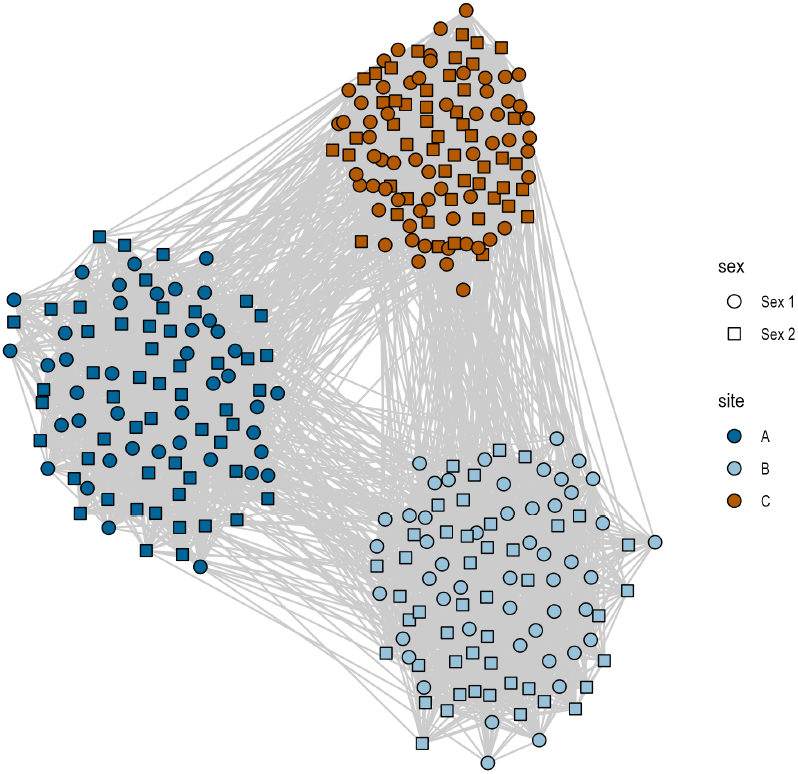
A single realisation of the differential site match model, ℳ_2_.

**Fig. 9.**
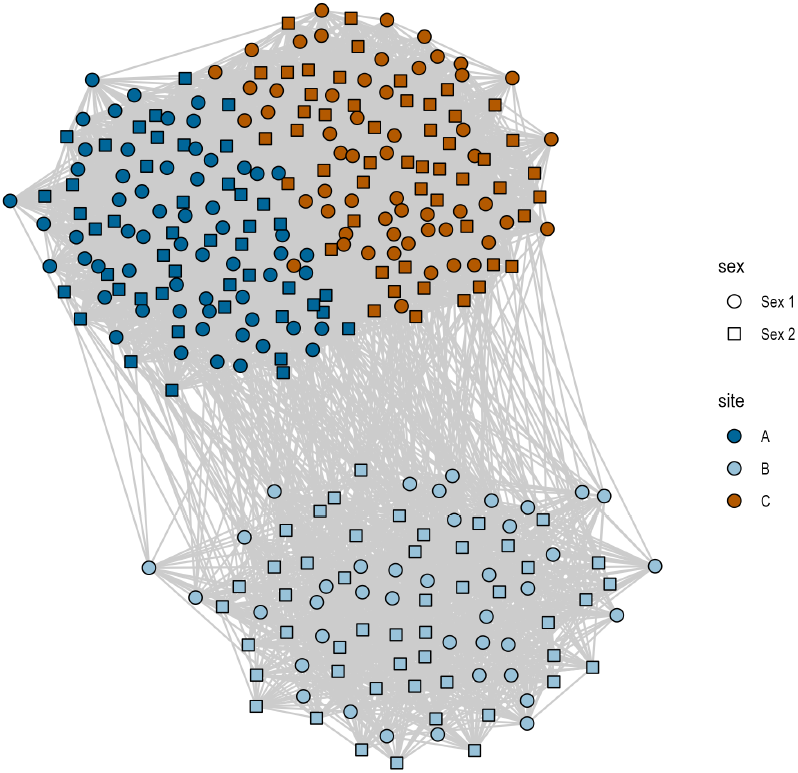
A single realisation of the site mix model, ℳ_3_.

Here, *θ*_1_ = *θ*_2_ = 3, corresponding to 14.95-fold increase in the probability of individuals being connected if they share the same burial site, and are buried at site A or B. However *θ*_3_ = 4, meaning that if both individuals are buried at C, then the probability is increased to 27.79-fold, compared to individuals buried at different sites.

#### B.3 The site mix model

ℳ_3_. The site mix model makes two individuals equally more likely to share a connection if they are buried at the same site, similar to model ℳ_1_. However, it also allows sites A and C to be more connected than sites, with no additional probability for connections between sites A and B, or sites B and C.

Here, *θ*_1_ = *θ*_2_ = *θ*_3_ = 3, corresponding to 14.95-fold increase in the probability of individuals being connected if they share the same burial site. However, there is still a 6.62-fold increase in the probability of a connection between individuals at sites A and C, resulting in a network in which sites A and C appear to “overlap”, which site B is more pronounced in its separation.

#### B.5 The sex match model

ℳ_4_. The sex match model makes two individuals equally more likely to share a connection if they are of the same genetic sex. This is similar to the site match model, and is included to test models where the variables has only two possible levels.

Here, *θ*_6_ = *θ*_7_ = 3, corresponding to 2.07-fold increase in the probability of individuals being connected if individuals share the same genetic sex. This is an unlikely scenario and is included for comparison to the three-level site match model.

**Fig. 10.**
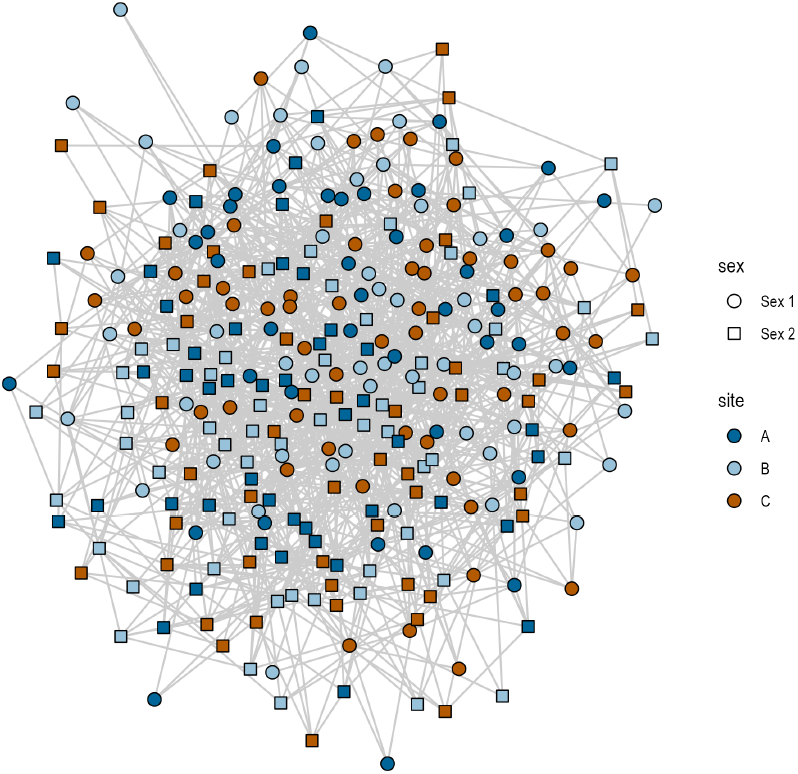
A single realisation of the sex match model, ℳ_4_.

#### B.6 The differential sex model

ℳ_5_. The differential sex model makes two individuals more likely to share a connection if they are of the same genetic sex, but this differs for different levels. If one of the coefficients were negative, this could be a result of matrilocality or patrilocality.

**Fig. 11.**
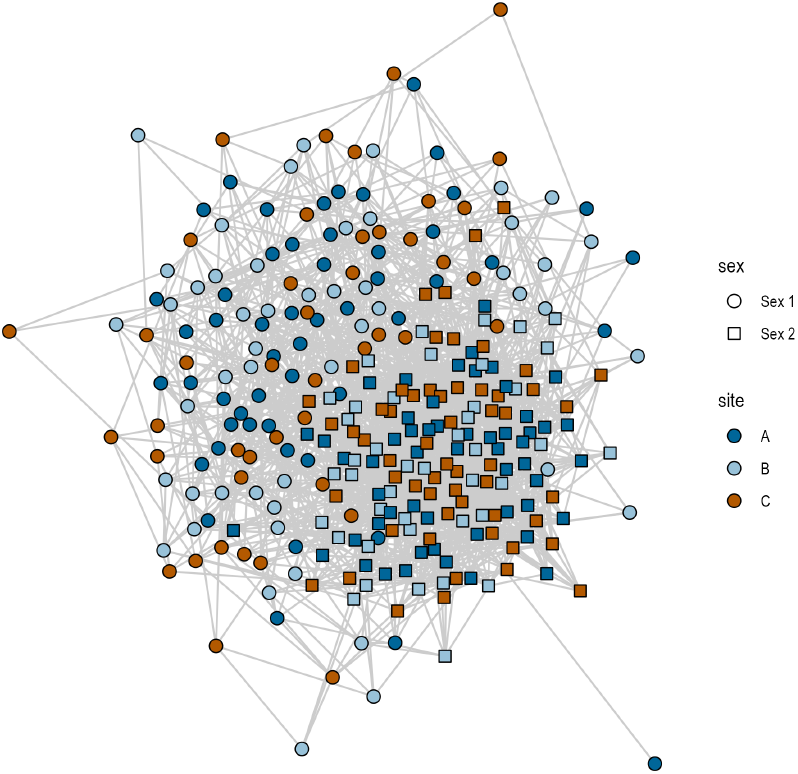
A single realisation of the differential sex match model, ℳ_5_.

In this case, if two individuals are both Sex 2, then they have they have a 4.21-fold increase in the probability of individuals being connected, but only a 2.07-fold increase if both are Sex 1.

#### B.7 The mixed homophily model

ℳ_6_. The mixed homophily makes both site location and genetic sex contribute to the probability of two individuals sharing a connection. The effect of sharing a site is greater than the effect of having the same genetic sex.

**Fig. 12.**
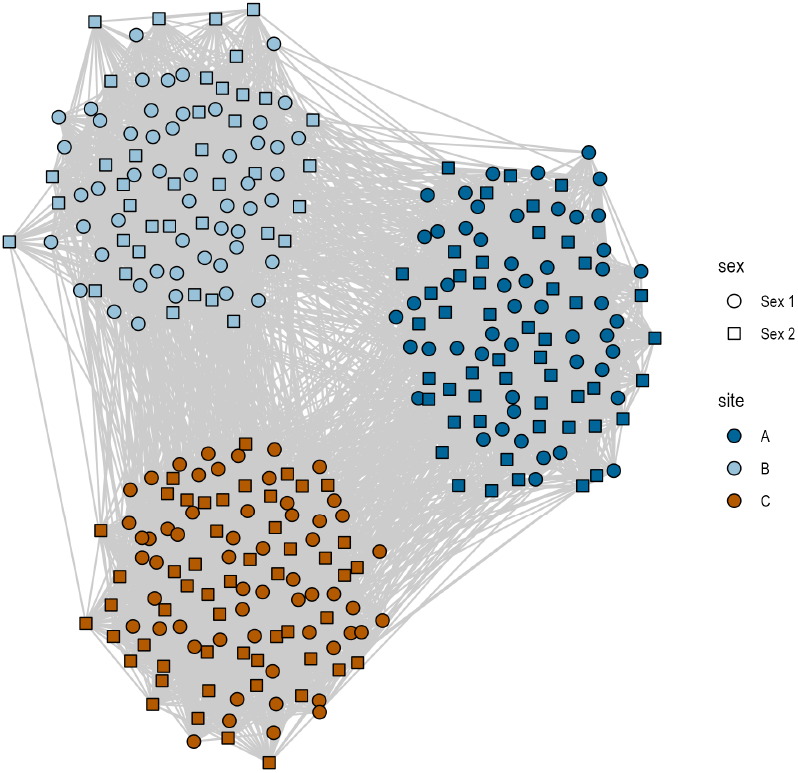
A single realisation of the mixed homophily model, ℳ_6_.

**Table 2.** Coefficient values of *θ*_*i*_ which differentiate the different models, ℳ_0_, …, ℳ_6_.

| Model ID | $\theta_0$ | $\theta_1$ | $\theta_2$ | $\theta_3$ | $\theta_4$ | $\theta_5$ | $\theta_6$ | $\theta_7$ |
| --- | --- | --- | --- | --- | --- | --- | --- | --- |
| $\mathcal{M}_0$ | -4 | 0 | 0 | 0 | 0 | 0 | 0 | 0 |
| $\mathcal{M}_1$ | -4 | 3 | 3 | 3 | 0 | 0 | 0 | 0 |
| $\mathcal{M}_2$ | -4 | 3 | 3 | 4 | 0 | 0 | 0 | 0 |
| $\mathcal{M}_3$ | -4 | 3 | 3 | 3 | 2 | 0 | 0 | 0 |
| $\mathcal{M}_4$ | -4 | 0 | 0 | 0 | 0 | 0 | 0.75 | 0.75 |
| $\mathcal{M}_5$ | -4 | 0 | 0 | 0 | 0 | 0 | 0.75 | 1.5 |
| $\mathcal{M}_6$ | -4 | 3 | 3 | 3 | 0 | 0 | 0.75 | 0.75 |

This model is the most complicated, but allows for the analysis of the effect of genetic sex, while still *accounting* for the effect of site. This might be of importance when there is a sampling bias at one site, say when yielding more individuals of Sex 1 at site A and more of Sex 2 at site B.

### C. Simulated model parameters

A concise table of model parameters are given below for the model equation differentiating the models from Supplementary Section B.

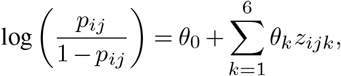

Note that we calculate the fold-change in probability from the coefficient values in the following way. For an coefficient value of *θ*^*1*^, and a baseline probability *θ*_0_ (the coefficient associated with “edges”) the fold-change, *φ*^*1*^ can be found via

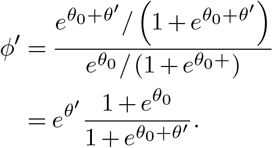

**Table 3.** Fold-increase values of *φ*_*i*_ (to two significant figures) which differentiate the different models, ℳ_0_, …, ℳ_6_.

| Model ID | $\phi_1$ | $\phi_2$ | $\phi_3$ | $\phi_4$ | $\phi_5$ | $\phi_6$ | $\phi_7$ |
| --- | --- | --- | --- | --- | --- | --- | --- |
| $\mathcal{M}_0$ | 0 | 0 | 0 | 0 | 0 | 0 | 0 |
| $\mathcal{M}_1$ | 15 | 15 | 15 | 0 | 0 | 0 | 0 |
| $\mathcal{M}_2$ | 15 | 15 | 28 | 0 | 0 | 0 | 0 |
| $\mathcal{M}_3$ | 15 | 15 | 15 | 6.6 | 0 | 0 | 0 |
| $\mathcal{M}_4$ | 0 | 0 | 0 | 0 | 0 | 2.1 | 2.1 |
| $\mathcal{M}_5$ | 0 | 0 | 0 | 0 | 0 | 2.1 | 4.2 |
| $\mathcal{M}_6$ | 15 | 15 | 15 | 0 | 0 | 2.1 | 2.1 |

### D. Model selection comparison

To compare the performance of model selection methods on the same data we calculated the Akaike Information criterion (AIC), the Bayesian Information criterion (BIC) for every realisation. Additionally, we attempted to use the p-values of coefficients in the model summary as a model selection method, retaining models where coefficients had p-value less than 0.05, and hence selection this model (called p-value method from here on). BIC outperformed both AIC and the p-value method as it was 100% accurate for all 700 realisations. AIC performed reasonably, but less well, with an overall accuracy of 94.71%. However, using AIC led to the null model being misclassified as models with either site or sex as significant variables in 16% of simulations, which would lead to a high false positive rate when no variables are significant.

**Fig. 13.**
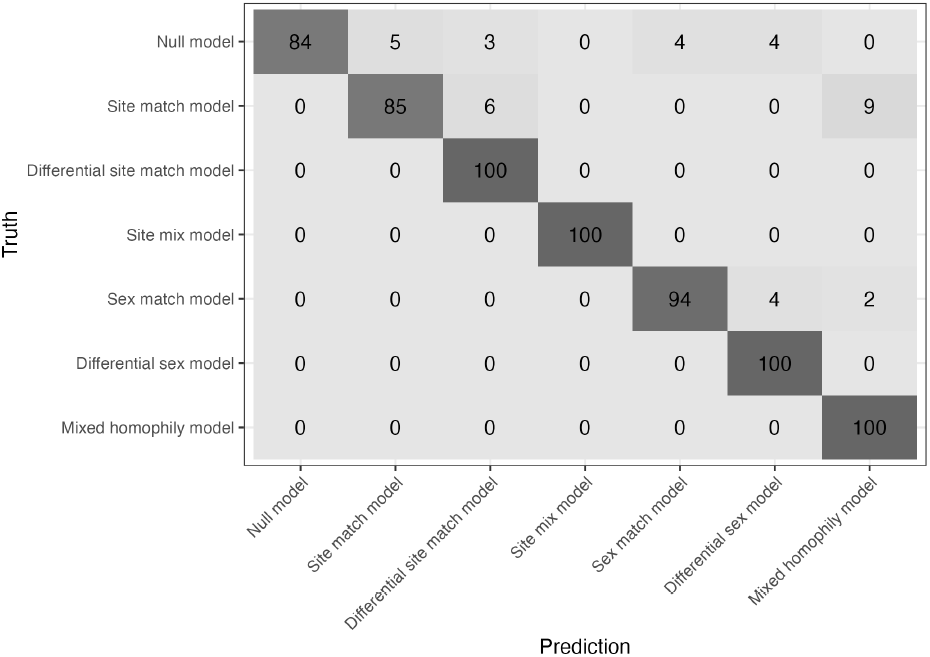
Confusion matrix for model selection accuracy using the Akaike Information criterion.

Conversely, the p-value method performed worse than either information criterion approach. The p-value method achieved only 47% accuracy, and suffered from issues of false positive classification for the null model (as well as one-parameter models), as well as clearly being biased towards more complicated, higher-dimensional models when the correct variables were identified.

**Fig. 14.**
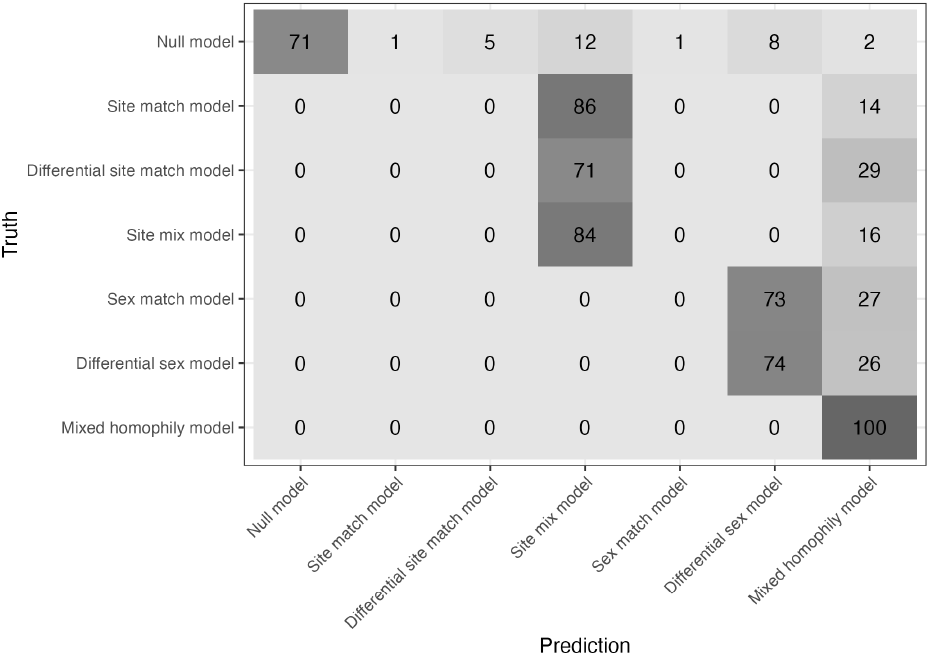
Confusion matrix for model selection accuracy using the p-value method.

Hence, based on these results, we suggest the use of BIC for model selection when using ERGMs.

### E. Empirical data

#### E.1 Data description

The empirical data in this study comes from three (exhaustively-sampled) Avar-associated sites in Hungary: Hajdúnánás-Fürj-halom-járás (HNJ, *n* = 11, 567-700 AD), Kunszállás-Fülöpjakab (KFJ, *n* = 37, 601-700 AD), Kunpeszér-Felsőpeszéri út (KUP, *n* = 13, 601-700 AD) and Rákóczifalva Bagi-földek 8 (RK, *n* = 176, 601-822 AD).The data was sequenced at the Max Planck Institute for Evolutionary Anthropology following standardized protocols designed in Ancient DNA Core facility(15). The sequence data was imputed using GLIMPSE(22) and IBD blocks were called using ancIBD(8). For a full description of how the data was processed, see Gnecchi-Ruscone *et al*.(15).

We defined individuals as “related” if they shared at least two blocks of IBD of length 12cM, and at least one block of IBD of length 16cM. The genetic sex of the individuals was estimated using the ratio of coverage on the X and Y chromosomes versus coverage on the autosomes(15).

For each individual we recorded the site at which they were found (“HNJ”, “KFJ”, “KUP” or “RK”), the period in which they lived (Early Avar period “EA” or Middle/Late Avar period “MA/LA”), the combined genetic sex and age of the individual (“XX/Adult”, “XX/Subadult”, “XY/Adult” or “XY/Subadult”), the burial orientation of the individual (“N-S”, “NNW-SSE” or “NW-SE”), whether the individual was buried with a seemingly random item in the form of an iron buckle (“Iron Buckle” or “No Iron Buckle”), and whether the individual was buried with a valuable item, potentially due to inherited status, in a golden item (“Gold” or “No Gold”).

#### E.2 Centrality measures

We began by clustering the vertices using the Louvain clustering algorithm (see Figure 15 A and B)(1). We observe that HNJ is separated into just two clusters, and that KUP and KFJ form two clusters each, with one being shared between them. Further, we see that RK forms nine clusters with three major clusters representing a large amount of within-site clustering that is not observed at the other sites.

**Fig. 15.**
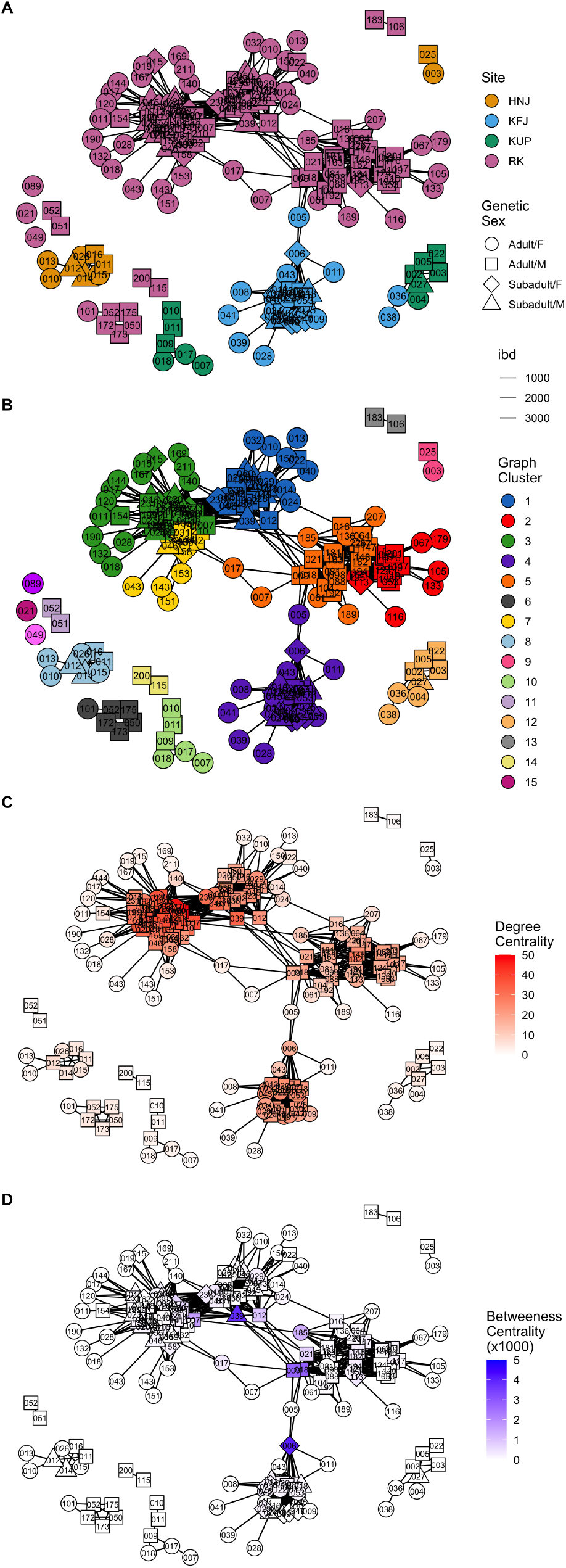
Network representations of the Avar data set with (A) site, (B) Louvain clustering, (C) degree centrality and (D) betweenness centrality indicated by vertex colours. Shape indicates the age and sex of the individuals.

We then inspected the connectivity in the model by calculating the degree centrality which measures the number of vertices each vertex is connected to (see Figure 15 C). We see that the main cluster at KFJ, and the three major sub-clusters at RK, indicate the highest amount of connectivity, indicating that these pedigrees and sites were simply larger.

Finally we looked at the betweenness centrality which measures vertices which are important to “connectedness” in the network (see Figure 15 D). A simple interpretation here would be high values of betweennes centrality indicate individuals who connect distinct, but connected, clusters. We find that the only obvious connecting individuals are within RK and KFJ, again connecting the four major clusters mentioned in the analysis of the degree centrality measures.

## Notes

### Competing Interest Statement

The authors have declared no competing interest.

## Bibliography

1. Pascal Held, Benjamin Krause, and Rudolf Kruse. Dynamic clustering in social networks using louvain and infomap method. In 2016 Third European Network Intelligence Conference (ENIC), pages 61–68. IEEE, 2016.

2. Gabor Csardi, Tamás Nepusz, Vincent Traag, Szabolcs Horvát, Fabio Zanini, Daniel Noom, and Kirill Müller. igraph: Network analysis and visualization in r. R package version, 1(1): 10–5281, 2023.

3. Chao Ning, Fan Zhang, Yanpeng Cao, Ling Qin, Mark J Hudson, Shizhu Gao, Pengcheng Ma, Wei Li, Shuzheng Zhu, Chunxia Li, et al. Ancient genome analyses shed light on kinship organization and mating practice of late neolithic society in china. Iscience, 24(11), 2021.

4. Jessica Pearson, Jane Evans, Angela Lamb, Douglas Baird, Ian Hodder, Arkadiusz Marciniak, Clark Spencer Larsen, Christopher J Knüsel, Scott D Haddow, Marin A Pilloud, et al. Mobility and kinship in the world’s first village societies. Proceedings of the National Academy of Sciences, 120(4):e2209480119, 2023.

5. Valéria Romano, Sergi Lozano, and Javier Fernández-López de Pablo. A multilevel analytical framework for studying cultural evolution in prehistoric hunter–gatherer societies. Biological Reviews, 95(4):1020–1035, 2020.

6. Linxuan Wang, Chen Duan, and Chao Ning. Genetic insights into ancient kinship and human history: Methods, applications, and implications. Nature Anthropology, 3(2):10009, 2025.

7. Adam B Rohrlach, Jonathan Tuke, Divyaratan Popli, and Wolfgang Haak. Breadr: An r package for the bayesian estimation of genetic relatedness from low-coverage genotype data. bioRxiv, pages 2023–04, 2023.

8. Harald Ringbauer, Yilei Huang, Ali Akbari, Swapan Mallick, Nick Patterson, and David Reich. ancibd-screening for identity by descent segments in human ancient dna. BioRxiv, 2023.

9. Divyaratan Popli, Stéphane Peyrégne, and Benjamin M Peter. Kin: a method to infer relatedness from low-coverage ancient dna. Genome Biology, 24(1):10, 2023.

10. Erkin Alaçamlı, Thijessen Naidoo, Merve N Güler, Ekin Sağ lıcan, Ş evval Aktürk, Igor Mapelli, Kıvılcım Başak Vural, Mehmet Somel, Helena Malmström, and Torsten Günther. Readv2: advanced and user-friendly detection of biological relatedness in archaeogenomics. Genome Biology, 25(1):216, 2024.

11. Sandra Penske, Mario Küßner, Adam B Rohrlach, Corina Knipper, Jan Nováček, Ainash Childebayeva, Johannes Krause, and Wolfgang Haak. Kinship practices at the early bronze age site of leubingen in central germany. Scientific Reports, 14(1):3871, 2024.

12. Maïté Rivollat Adam Benjamin Rohrlach, Harald Ringbauer, Ainash Childebayeva, Fanny Mendisco, Rodrigo Barquera, András Szolek, Mélie Le Roy, Heidi Colleran, Jonathan Tuke, et al. Extensive pedigrees reveal the social organization of a neolithic community. Nature, 620(7974):600–606, 2023.

13. Vanessa Villalba-Mouco, Camila Oliart, Cristina Rihuete-Herrada, Adam B Rohrlach, María Inés Fregeiro, Ainash Childebayeva, Harald Ringbauer, Iñigo Olalde, Eva Celdrán Beltrán, Catherine Puello-Mora, et al. Kinship practices in the early state el argar society from bronze age iberia. Scientific Reports, 12(1):22415, 2022.

14. Chris Fowler, Inigo Olalde, Vicki Cummings, Ian Armit, Lindsey Büster, Sarah Cuthbert, Nadin Rohland, Olivia Cheronet, Ron Pinhasi, and David Reich. A high-resolution picture of kinship practices in an early neolithic tomb. Nature, 601(7894):584–587, 2022.

15. Guido Alberto Gnecchi-Ruscone, Zsófia Rácz, Levente Samu, Tamás Szeniczey, Norbert Faragó, Corina Knipper, Ronny Friedrich, Denisa Zlámalová, Luca Traverso, Salvatore Liccardo, et al. Network of large pedigrees reveals social practices of avar communities. Nature, 629(8011):376–383, 2024.

16. Ke Wang, Bendeguz Tobias, Doris Pany-Kucera, Margit Berner, Sabine Eggers, Guido Alberto Gnecchi-Ruscone, Denisa Zlámalová, Joscha Gretzinger, Pavlína Ingrová, Adam B Rohrlach, et al. Ancient dna reveals reproductive barrier despite shared avar-period culture. Nature, pages 1–8, 2025.

17. Michael Schweinberger, Pavel N Krivitsky, Carter T Butts, and Jonathan R Stewart. Exponential-family models of random graphs. Statistical Science, 35(4):627–662, 2020.

18. S Golshid Sharifnia and Abbas Saghaei. A statistical approach for social network change detection: an ergm based framework. Communications in Statistics-Theory and Methods, 51(7):2259–2280, 2022.

19. Gideon Schwarz. Estimating the dimension of a model. The annals of statistics, pages 461–464, 1978.

20. Mathias Drton and Martyn Plummer. A bayesian information criterion for singular models. Journal of the Royal Statistical Society Series B: Statistical Methodology, 79(2):323–380, 2017.

21. Jacob Cohen. A power primer. 2016.

22. Simone Rubinacci, Diogo M Ribeiro, Robin J Hofmeister, and Olivier Delaneau. Efficient phasing and imputation of low-coverage sequencing data using large reference panels. Nature genetics, 53(1):120–126, 2021.

